# Identifying Artifacts from Large Library Docking

**DOI:** 10.1101/2024.07.17.603966

**Authors:** Yujin Wu, Fangyu Liu, Isabella Glenn, Karla Fonseca-Valencia, Lu Paris, Yuyue Xiong, Steven V. Jerome, Charles L. Brooks, Brian K. Shoichet

## Abstract

While large library docking has discovered potent ligands for multiple targets, as the libraries have grown, the very top of the hit-lists can become populated with artifacts that cheat our scoring functions. Though these cheating molecules are rare, they become ever-more dominant with library growth. Here, we investigate rescoring top-ranked molecules from docking screens with orthogonal methods to identify these artifacts, exploring implicit solvent models and absolute binding free energy perturbation (AB-FEP) as cross-filters. In retrospective studies, this approach deprioritized high-ranking non-binders for nine targets while leaving true ligands relatively unaffected. We tested the method prospectively against results from large library docking AmpC β-lactamase. From the very top of the docking hit lists, we prioritized 128 molecules for synthesis and experimental testing, a mixture of 39 molecules that rescoring flagged as likely cheaters and another 89 that were plausible true actives. None of the 39 predicted cheating compounds inhibited AmpC up to 200µM in enzyme assays, while 57% of the 89 plausible true actives did do so, with 19 of them inhibiting the enzyme with apparent K_i_ values better than 50µM. As our libraries continue to grow, a strategy of catching docking artifacts by rescoring with orthogonal methods may find wide use in the field.

**Graphical TOC Entry:** 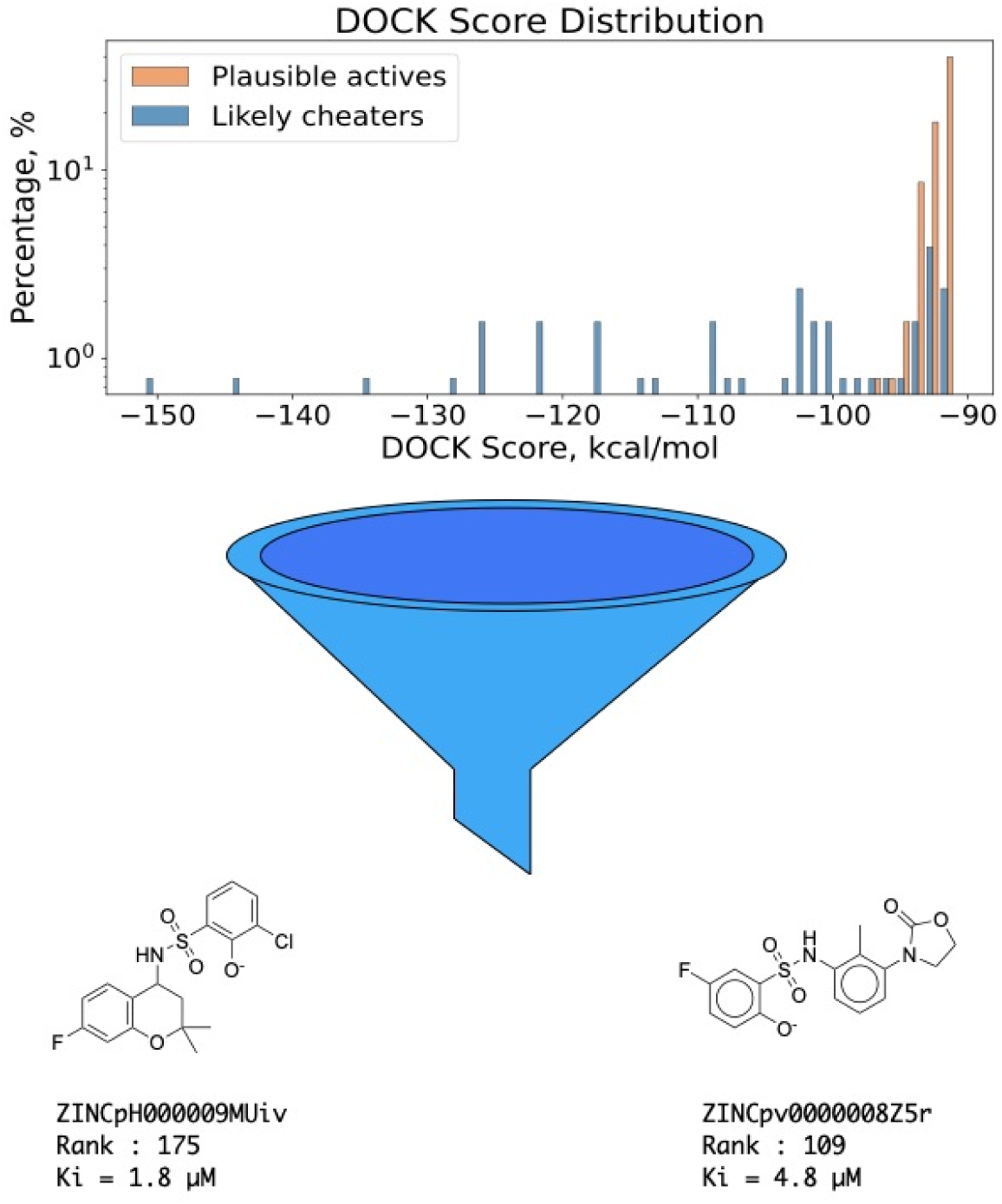

## Introduction

In the last five years, the number of readily available molecules for ligand discovery has grown from several million to tens of billions. Structure-based docking of those new libraries has revealed new chemotypes with potent affinity for targets ranging from enzymes [1–3], to GPCRS [4–11], to transporters [5, 12], to kinases [6, 13]. As the libraries have grown, however, both simulations and experiment have shown that the very top of the docking- ranked list becomes populated with molecules that "cheat" the scoring function. These false positives can be classified into normal docking failures and into artifacts (Figure 1). Normal docking failures, while common, occur through well-known problems of balancing different energy terms in the scoring functions. They often inhabit the same chemical space as true ligands and spread relatively evenly through the docking results. What we call cheating molecules, conversely, are rare molecules that through failures of parameterization or adoption of unusual structures rank among the very top molecules, often with scores off-set from those of the overall distribution of docked molecules. As our virtual libraries continue to grow [8, 14–19], such cheating molecules may have ever greater impact on virtual screening hit lists.

**Figure 1:**
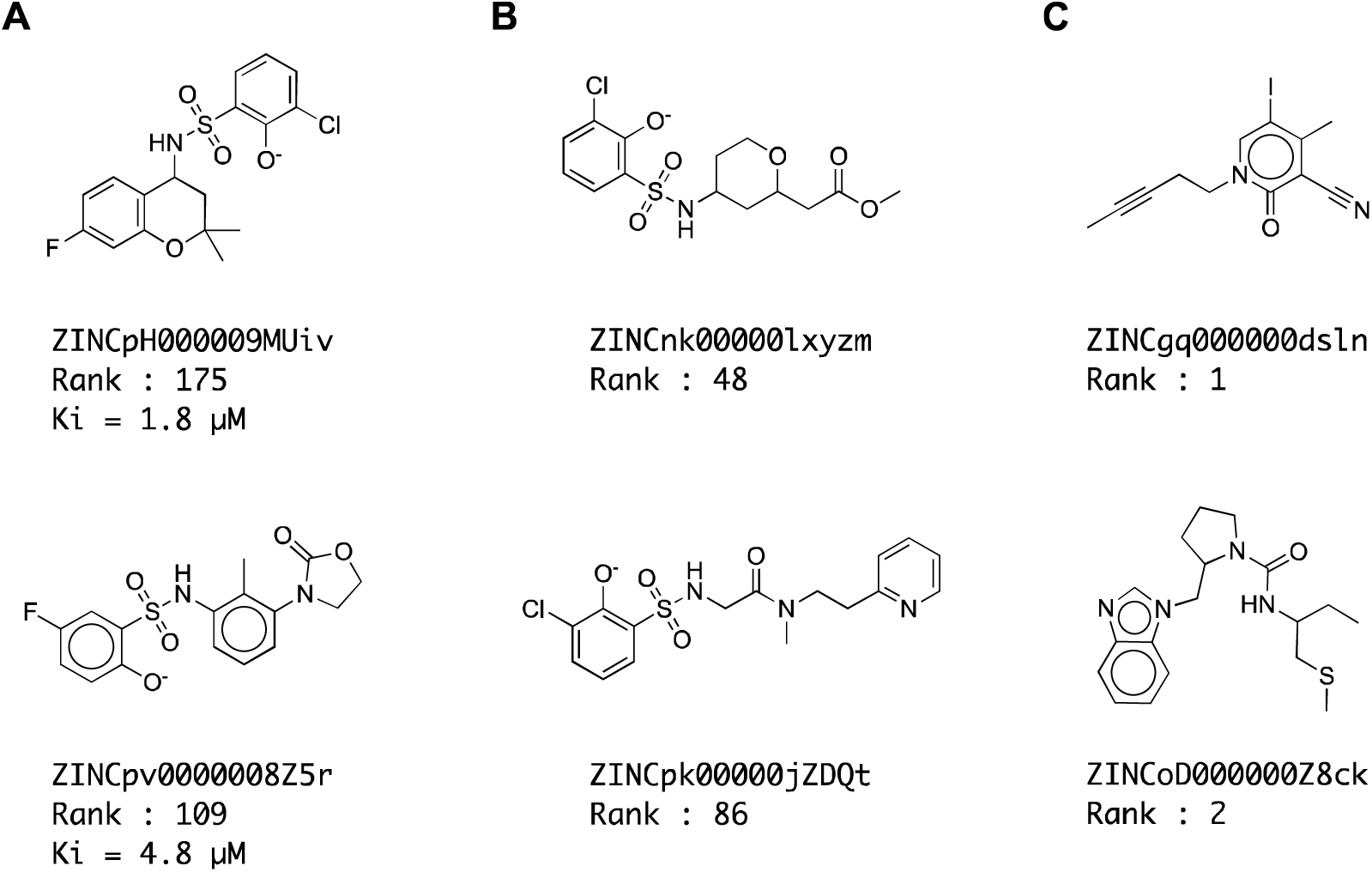
Examples of (A) true inhibitors, (B) normal docking failures, and (C) cheating compound against AmpC. Ranks are out of 1.7 billion docked.

Since these cheaters arise from holes in a particular scoring function, rescoring high- ranking docked molecules with a second function may identify them as outliers. Here, we cross-filter high-ranking docked molecules with three different methods: we compare DOCK3.8 implicit solvation energies to those calculated by FACTS (Fast Analytical Continuum Treatment of Solvation) [20] and to those calculated by GBMV (Generalized Born using Molecular Volume) [21], and compare DOCK3.8 rankings to those from an AB-FEP (Absolute Binding-Free Energy Perturbation) calculation on the top-ranking molecules. These calculations are undertaken both retrospectively and prospectively, where 39 cheating artifacts identified by rescoring are synthesized and tested experimentally, as are 89 molecules judged to be likely ligands by the same rescoring strategy. The results of these experiments suggest that a rescoring approach may be useful to deprioritize cheating artifacts that can concentrate among the top-ranking molecules from large library docking.

## RESULTS

Retrospective study on a target with hundreds of experimental ligands and decoys. A good system to retrospectively optimize the cross-filtering approach is one where large libraries of molecules are docked and where hundreds of docking false- and true-positives are measured by experiment. There are only a handful of these, with among the richest sets being against the enzyme AmpC β-lactamase. In a recent study, 1.71 billion molecules were docked against AmpC, and over 1400 molecules tested experimentally [Liu, 2024]. As in earlier studies, the distribution of docking scores from this campaign suggested that cheating molecules might be present (Figure 2). As the library increased in size, there was a regular improvement in docking scores, until around 300 million molecules, when molecules began to appear whose very favorable docking scores diverged from the rest of the distribution. As the library grew further, more of these molecules appeared, reaching ever better scores. Both simulation [5] and experience with other systems [1] suggested that these might be cheating molecules.

**Figure 2:**
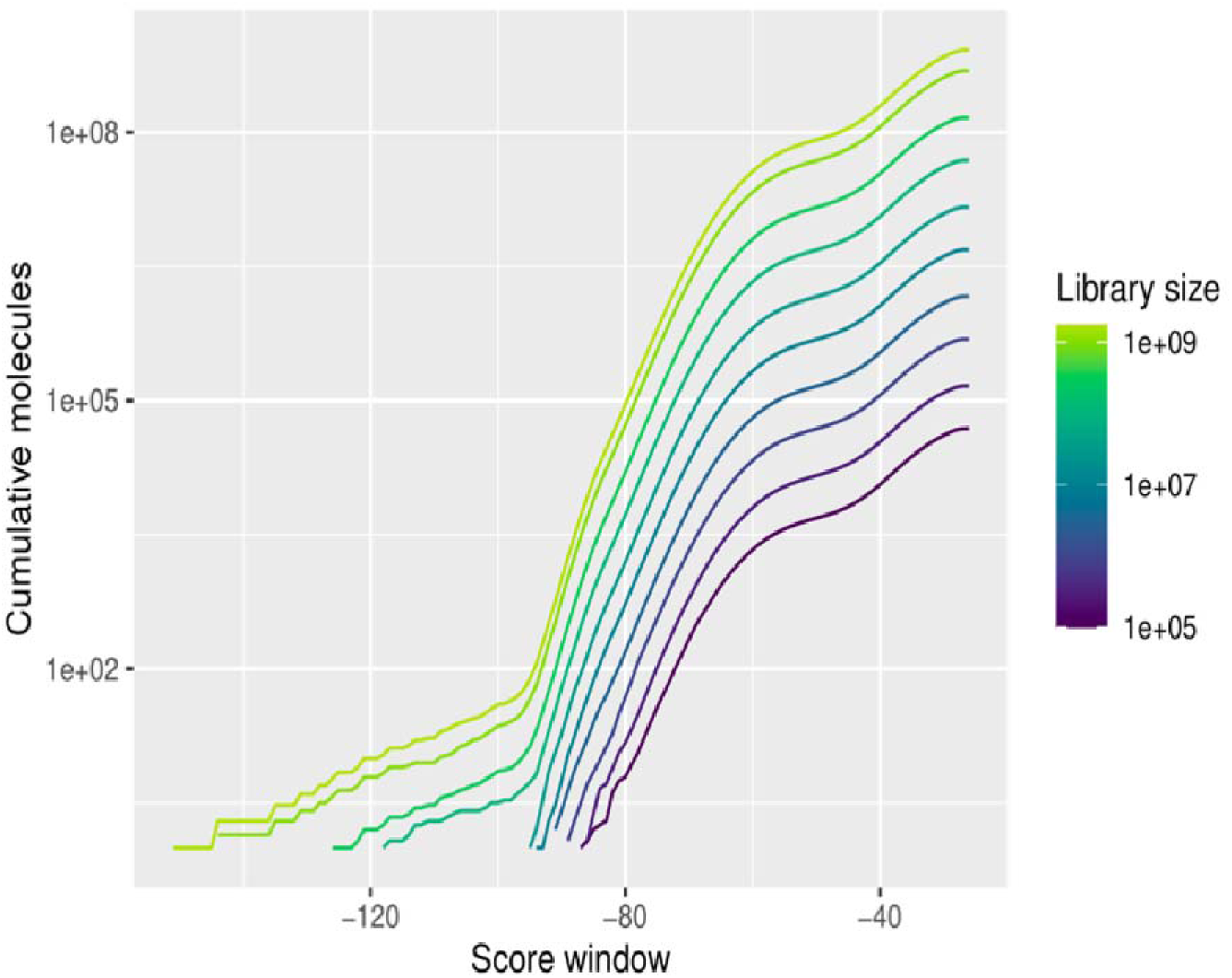
The distribution of docking scores with library growth against AmpC. As the library climbs towards a billion molecules, docking hits with unusually favorable (more negative) scores begin to appear.

We set out to test the cross-filtering approach on the 1439 docked-and-experimentally tested AmpC hits, asking whether it could separate true ligands from non-binders while minimizing false-negatives. We began by calculating FACTS solvation energies for the 1439 compounds, which took an average of 90 seconds per molecule on a Linux CPU. Plotting the normalized DOCK3.8 and FACTS desolvation energies against each other revealed a bivariant normal distribution (Figure 3A). Clustering those molecules within 3σ of the mean of both distributions reveals an ellipse outside of which 268 of the molecules fall; these molecules are flagged as having unusual solvation energies (i.e., outliers of the solvation energy distribution) by one of methods. Of these, 262 do not bind AmpC detectably up to 200µM, while six were decent inhibitors, the best of which had an affinity of 9.7µM. Thus, insisting that all molecules are within 3σ of both removed more than 22% of the non-binders while only losing 3.8% of the true ligands, none of which was among the most potent (which ranged down to sub-µM K_d_ values). While we’d prefer not to lose any of the true ligands, we do note that this was a stringent test of the method, as it is applied to all the molecules tested in this campaign, not only the thin wedge of unusually well-scoring molecules where we expect the “cheaters” to concentrate.

**Figure 3:**
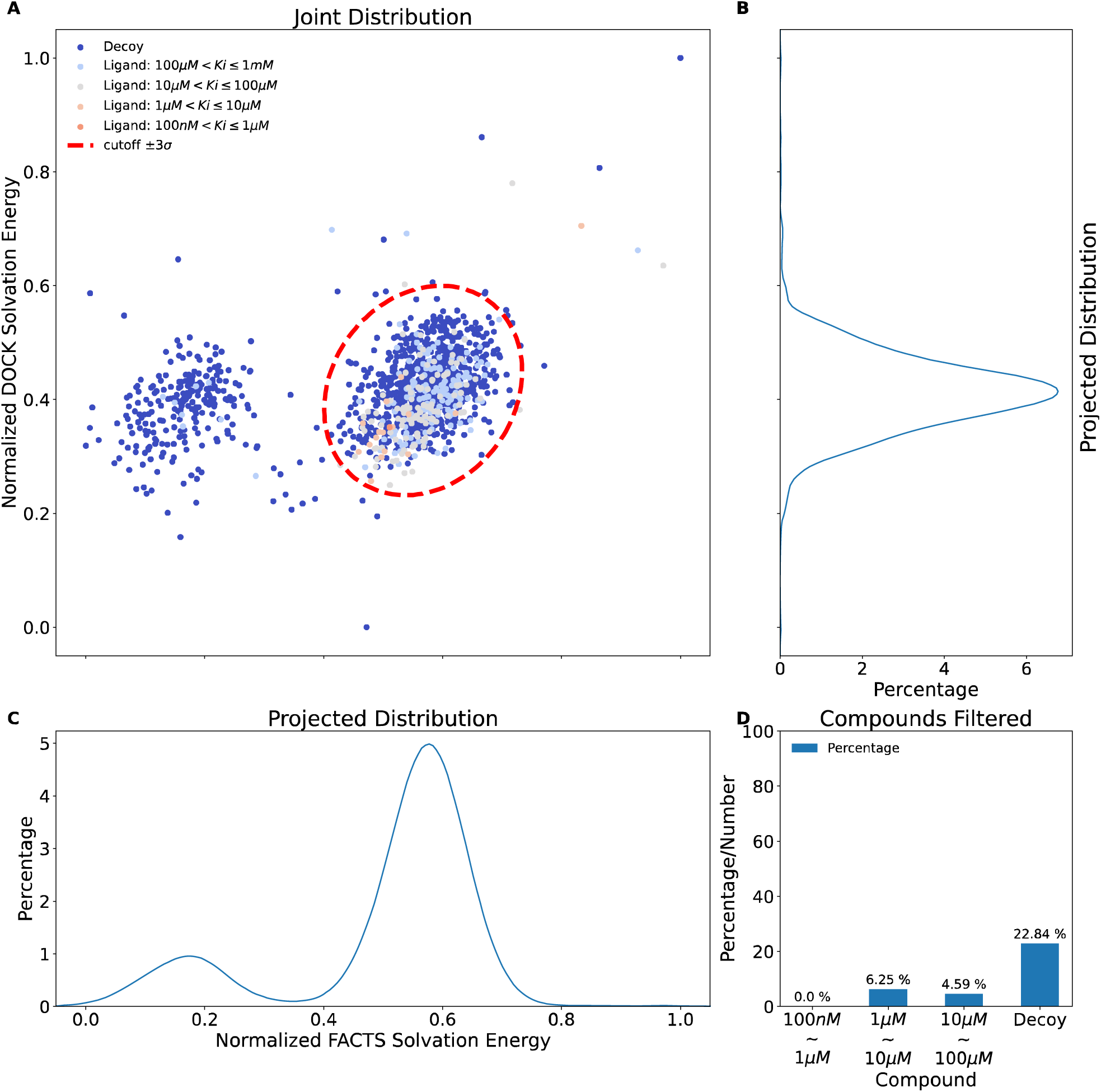
FACTS rescoring results for 1,440 experimentally validated compounds against AmpC. (A) Joint distribution of normalized DOCK and FACTS solvation free energy contribution. Distribution of (B) DOCK and (C) FACTS solvation free energy contribution. (D) Percentage of ligands and decoys filtered.

Encouraged by these results, we applied the same protocol against the σ2 and D4 receptors, which had the advantage of having had about 500 docking hits synthesized and experimentally tested against them [1, 6]; in this sense, they were akin to the AmpC screen, though at a smaller scale. Here too, cross-filtering successfully flagged 22% and 28% non-binders, respectively, (Figure 4), while only losing four and three true ligands, respectively, none of which were among the more potent found in these docking campaigns. Many of the filtered compounds corresponded to molecules that had unusually favorable docking scores and unusual physical features for a hit and might thus be cheating molecules – a point to which we will return in the prospective testing part of this study.

**Figure 4:**
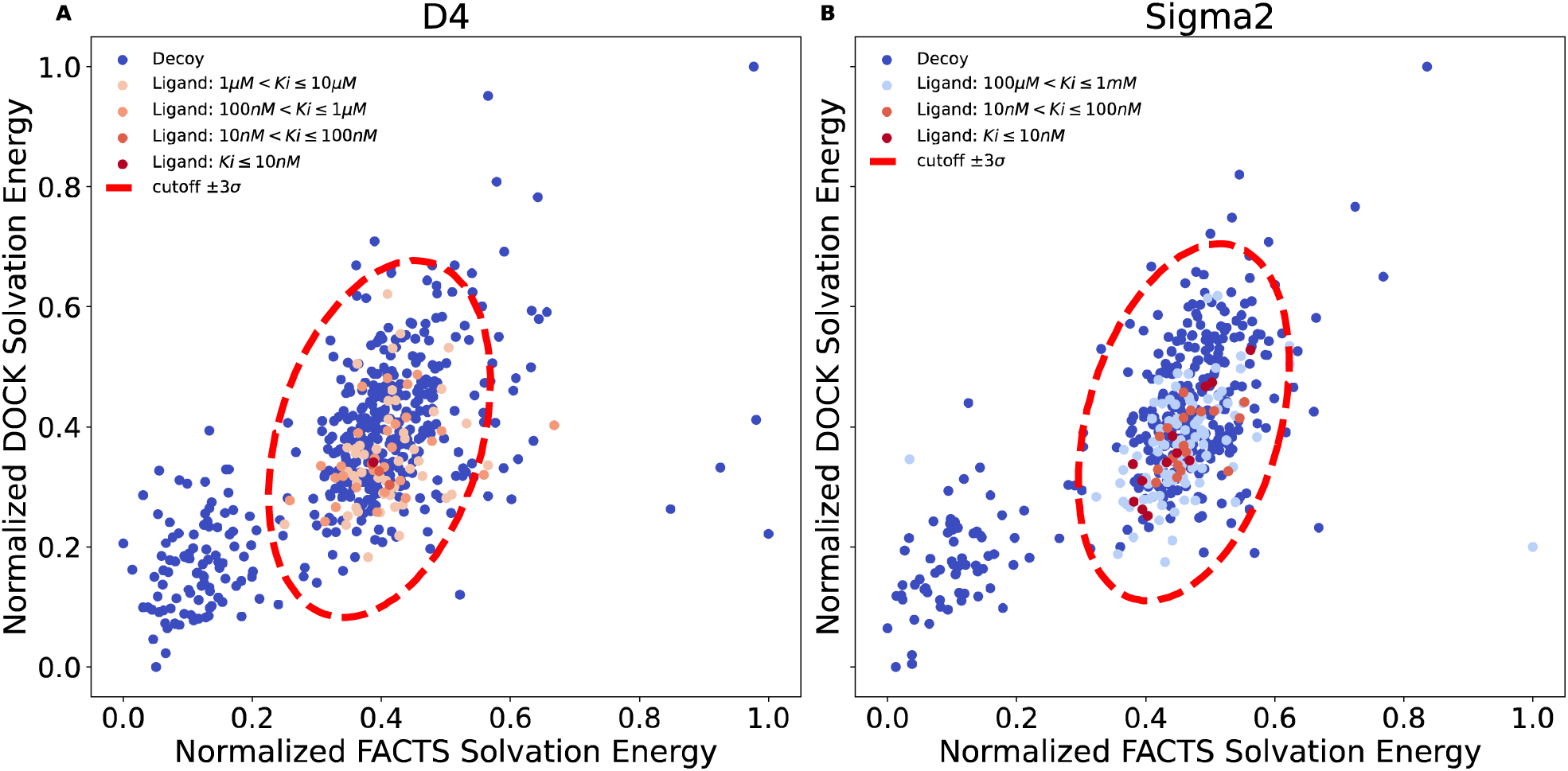
FACTS rescoring results for (A) D4 and (B) Sigma2. Total number of experimentally validated compounds are 537 and 495 respectively.

Cross filtering against nine receptors with different methods and parameters. To further investigate the robustness of this cross-filtering approach, we tested it against other docking targets with different binding pocket environments than those of AmpC, σ2, and D4. With the initial three studies establishing that this approach might be sensible, targeting a wider set of systems also afforded us the chance to try different rescoring methods, and to vary the parameters with which we did so (Table 1).

**Table 1.** Receptor Dataset.

| Receptor Name (Abbreviation) | Receptor Class | Docking hits<br>rescored |
| --- | --- | --- |
| AmpC β-lactamases (AmpC) | hydrolases | 300,000 |
| Serotonin transporter (SERT) [12] | transporter | 300,000 |
| Sigma-2 recetpor (Sigma2) [1] | membrane protein | 300,000 |
| SARS-CoV-2 macrodomain enzyme (Mac1) [22] | MAR-hydrolase | 500,000 |
| SARS-CoV-2 main protease (Mpro) [23] | cysteine hydrolase | 500,000 |
| Melation receptor type 1A (MT1) [11] | GPCR | 300,000 |
| Cannabinoid receptor type 1 (CB1) [24] | GPCR | 300,000 |
| Dopamine receptor D<sub>4</sub> (D4) [6] | GPCR | 300,000 |
| $\alpha 2a$ adrenergic receptor (Alpha2a) [9] | GPCR | 165,000 |

Here again, we sought cases where docked ligands had been tested, admittedly in the 40-molecule range rather than the 500 to 1500 as with σ2, dopamine D4, and AmpC, but nevertheless revealing both true ligands and false positives. There are by now close to twenty of theses [1, 4, 6, 8, 9, 11–13, 22–26], we focused on nine to which we had ready access (Table 1). Against each target hundreds of millions to billions of compounds had been docked; we rescored the several hundred-thousand top-ranking poses. For each of the nine targets, we explored rescoring not only with FACTS, as in the initial studies above, but also with a second implicit solvent method, GBMV, and explored several different calculation parameters (Table S1). As before, we note that this is not the true use case we envision for rescoring, as it is applied to all docked-and-tested compounds, not only those very top ranking ones where we might expect the cheaters to concentrate. Nevertheless, it provides a useful sanity check for the strategy.

We observed similar joint distribution for all nine targets using FACTS (Table 2, Figure 5). Of the 300,000 to 500,000 high-ranking compounds rescored, between 7 and 19% were outliers by cross-filtering, sitting outside the bounds of the 3σ score distribution. For six of the nine targets, no potent ligands were among these outliers while experimental non-binders were captured as outliers, representing 3.85 to 13.51% of the total docking false positives across the screens (Table 2). For three targets, Alpha2a adrenergic receptor, AmpC, Sigma2, one potent ligand was categorized as an outlier, representing 12.5, 4.0 and 3.7% of the total ligands found for these targets, respectively; none are among the strongest binders found for their receptors. The best performance was for the CB1 and MT1 receptors, and for SERT, where none of the true ligands are removed whatsoever. In these studies, we used FACTS with 1000 steps of minimization and ParamChem force field parameters; the joint distributions for each of the other three parameter setups are plotted in Supplementary Figures S1, S2, and S3, where similar distributions were observed. This suggests that the approach is robust to different parameters and to different implicit solvent models.

**Figure 5:**
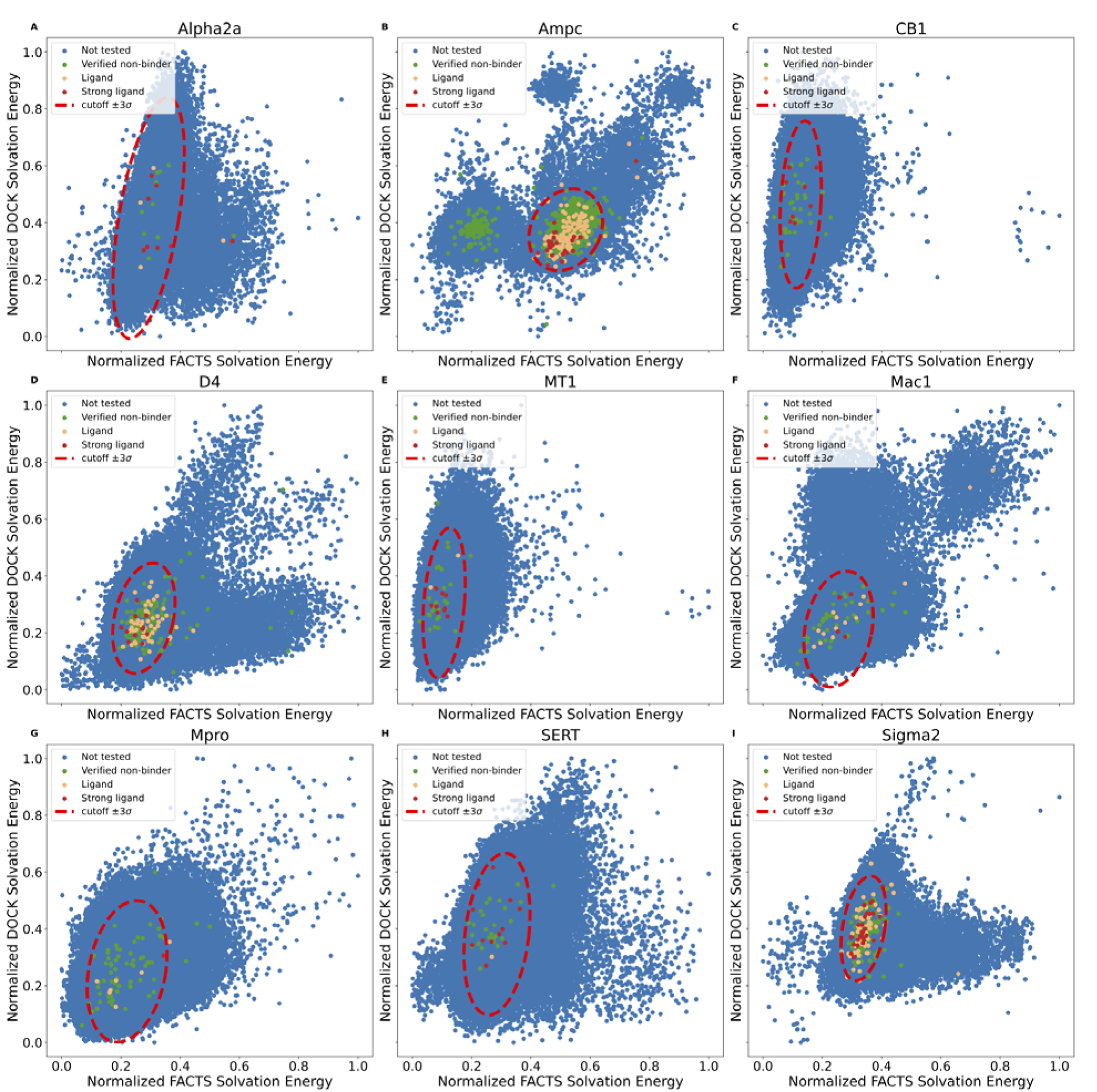
The joint distribution of the solvation free energy for receptor target (A) Alpha2a, (B) AmpC, (C) CB1, (D) D4, (E) MT1, (F) Mac1, (G) Mpro, (H) SERT and (I) Sigma2. Relatively potent ligands are colored red, and the rest of ligands are colored in orange. Experimentally tested non-binders are colored green and compounds that were not tested are colored blue.

**Table 2:** Number/Percentage of Compounds Removed by Cross Filtering.

| Receptor | Ki cutoff ($\mu M$)<sup>a</sup> | Potent Ligand | Decoy<sup>b</sup> | Compounds<sup>c</sup> |
| --- | --- | --- | --- | --- |
| Alpha2a | 3.0 | 1 / 12.5% | 1 / 10.00% | 6239 / 7.29% |
| AmpC | 10.0 | 1 / 4.0% | 124 / 16.56% | 55592 / 18.82% |
| CB1 | 10.0 | 0 / 0.0% | 5 / 13.51% | 34197 / 11.65% |
| D4 | 0.3 | 0 / 0.0% | 18 / 14.29% | 24411 / 8.34% |
| MT1 | 3.0 | 0 / 0.0% | 3 / 11.54% | 30934 / 10.55% |
| Mac1 | 300.0 | 0 / 0.0% | 8 / 19.51% | 26954 / 9.35% |
| Mpro | 100.0 | 0 / 0.0% | 6 / 8.70% | 31276 / 10.83% |
| SERT | 10.0 | 0 / 0.0% | 1 / 3.85% | 29788 / 10.17% |
| Sigma2 | 0.3 | 1 / 3.7% | 10 / 9.09% | 20734 / 7.08% |

<sup>a</sup> The upper limit of the binding affinity for potent ligands. <sup>b</sup> Experimentally validated non-binders. <sup>c</sup> All high ranking compounds.

More compounds are flagged as cheaters among top ranking results. The true use case for cross-filtering is to remove molecules that cheat the scoring function, which experience (Figure 2) and simulation [5] suggest occur among the very highest-ranking compounds. It is thus interesting to understand how the number of docking hits captured as outliers changes with docking rank. Overall, cross-filtering highlighted about 10% of the high-ranking docking hits from among the top 300,000 to 500,000 molecules (Table 2). If we plot the number of outliers captured by the cross filtering across the rank distribution, however, we find higher fractions of outliers among the very top-ranking compounds for six of the nine targets (SI Figure 4).

Prospective Study – Are Cheaters Experimental Non-Binders? To test cross-filtering prospectively, we prioritized and had synthesized the very top-ranking 128 de novo molecules from the AmpC screen [Liu, 2024] (ranked 1 to 415 out of 1.7 billion, DOCK3.8 scores from -151 to -91 kcal/mol). None of these had been previously tested. Of these, solvation cross-filtering identified 39 as cheaters (ranked 1 to 407) falling outside of the 3σ radii of the mean scores of the joint distribution (Figure 6a), and 89 as non-cheaters falling within that distribution. Given their good scores, these 89 were considered plausible AmpC inhibitors; their scores overlapped with those of the cheaters (Figure 6b).

**Figure 6:**
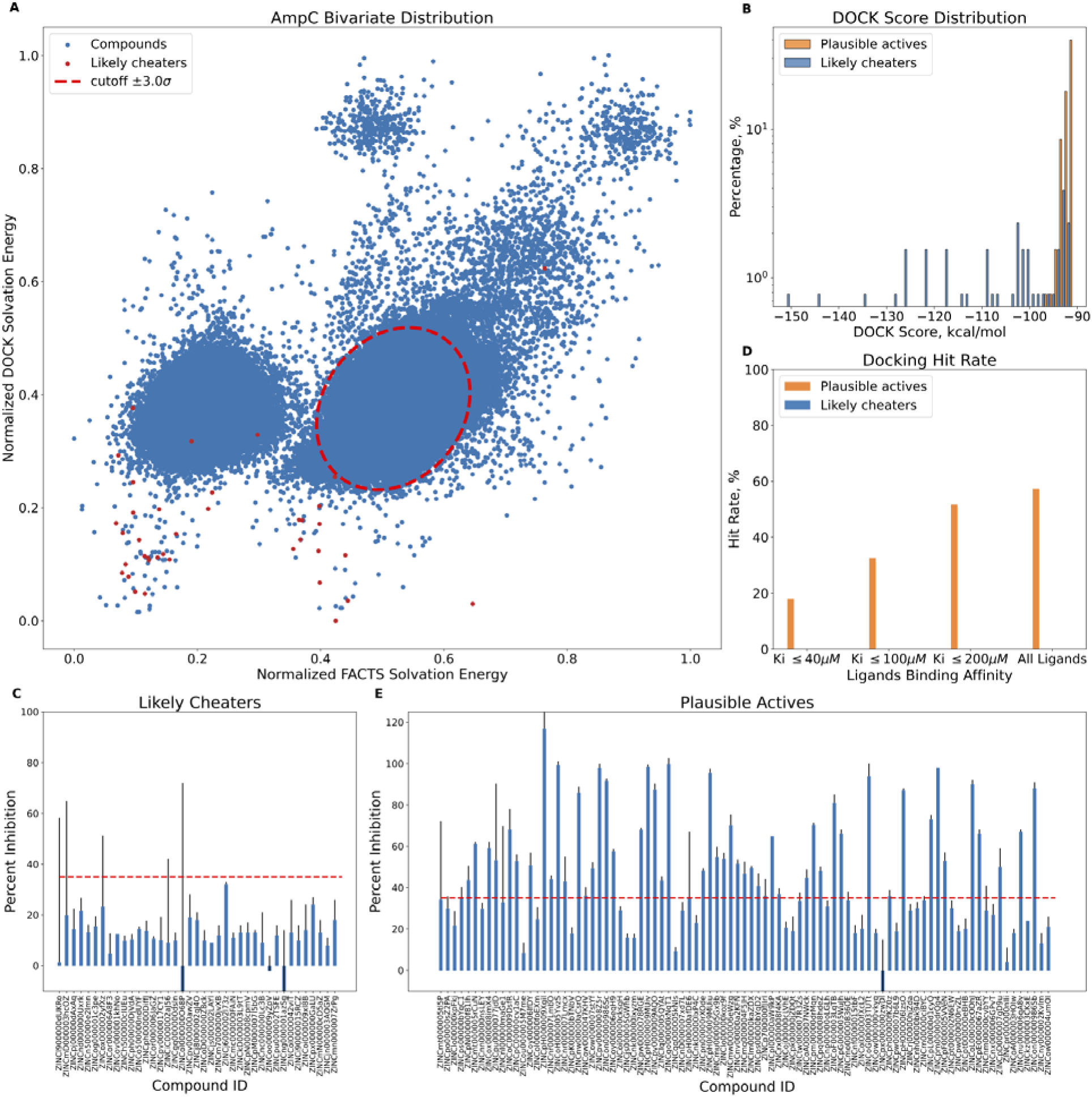
Experimental testing of predicted high-ranking docking cheaters and plausible true actives against AmpC. (A) Bivariate distribution of AmpC top 300K ranking compounds. The predicted cheaters are highlighted in red. (B) DOCK score distribution for top ranking putative cheaters and plausible true actives. (C) Percentage inhibition at 200 µM for likely cheaters (D) Docking hit rate as a function of AmpC apparent . (E) Percentage inhibition at 200 µM for plausible actives.

All 128 molecules were tested for AmpC inhibition, initially at 200, 100, and 40 µM. None of the 39 outliers inhibited the enzyme substantially at even the top concentration, including compounds ranked 1 to 26 of the 1.7 billion docked; all were classified as non-binders (Figure 6c). Conversely, of the 89 high-ranking plausible ligands, 51 (57%) inhibited meaningfully at the top concentration (Figure 6d), with inhibition at the lower concentrations consistent with apparent Ki values of 200 µM or better, and with 19 inhibiting the enzyme with apparent K_i_ values between 1.8 and 50 µM (Figure 6e).

Concentration-response curves for the eight most potent inhibitors were well-behaved (Figure 7). Their activities place them among the more potent inhibitors discovered for AmpC from docking or high-throughput screens [6, 27, 28] [Liu, 2024] (full compound results are listed in ampc-prospective-result.xlsx), while the gross hit rate of 57% (Figure 6d) is among the highest observed for this enzyme. The experimental results and corresponding SMILES string for each compound are recorded in the Supplementary file (ampc-prospective- result.xlsx).

**Figure 7:**
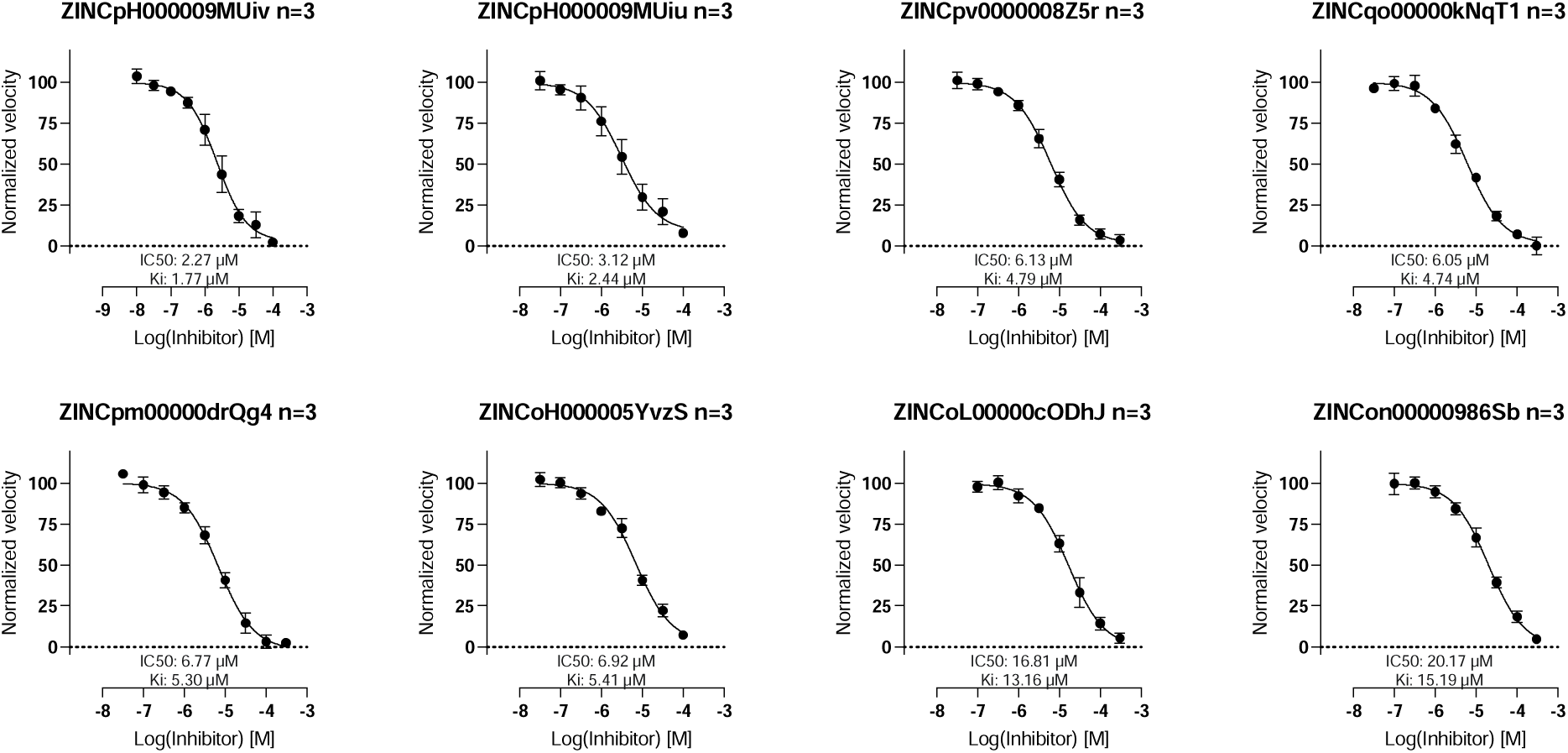
Concentration-response curves for top eight docking hits.

The AmpC assays were run in 0.01% Triton X-100, reducing the likelihood of colloidal aggregation. Nevertheless, for the 10 most potent inhibitors, dynamic light scattering (DLS) and counter-screening against Malate Dehydrogenase were used to investigate colloidal aggregation at concentrations 10-fold higher their apparent AmpC K_i_ values (Supplementary Figure S5 and S6). Only one compound, with apparent K_i_ of 8.6 µM, formed colloid-like particles, while none were observed for other top hits, consistent with these latter molecules acting as classic, active-site-directed inhibitors.

Other methods to detect cheaters – AB-FEP and MM/GBMV. In principle, one should be able to use many orthogonal scoring functions to identify molecules that cheat docking scoring functions. We tried two more here, absolute binding-free energy perturbation (AB-FEP) and molecular mechanics/Generalized Born Molecular Volume (MM/GBMV). Both represent higher levels of theory than implicit solvation methods like FACTS, especially AB-FEP. Both were prospectively applied to the same 128 top-ranking molecules described in the previous section (see Methods).

While GBMV with only minimization had performed relatively well in retrospective calculations (SI Table 1), the addition of a molecular dynamics component seemed to diminish its ability to prospectively distinguish the 89 plausible inhibitors or the 51 true ones, from the 39 cheating artifacts. A challenge here was knowing where to draw the cut- off between plausible inhibitors and likely cheating artifacts, since the MM/GBMV energies were much higher in magnitude than the energies inferred from the experimental apparent Ki values (Figure 8a). Accordingly, we investigated different energy cutoffs to distinguish the cheaters from the plausible ligands. Irrespective of where we drew the boundary, however, either too few true cheaters were found (true negatives), or too many true inhibitors were (false negatives). For instance, when we set the cutoff to be worse (greater) than -75 kcal/mol, only 13 of the 39 cheaters were identified, as was one phenolate sulfonamide, a close analog of true inhibitors and a plausible ligand (Figure 8b). As we raised the stringency of the cutoff, more cheaters were found but so were more true inhibitors. Increasing the cutoff to -125 kcal/mol identified 24 of the true cheaters but at the cost of adding 13 of the 89 plausible ligands, of which 9 were ultimately shown to be true inhibitors by experiment (Figure 8c) Thus, the higher sampling of MM/GBMV led to worse results than the simpler FACTS approach. This may reflect the increased noise on addition of the molecular dynamics and the large magnitude of the MM/GBMV energies.

**Figure 8:**
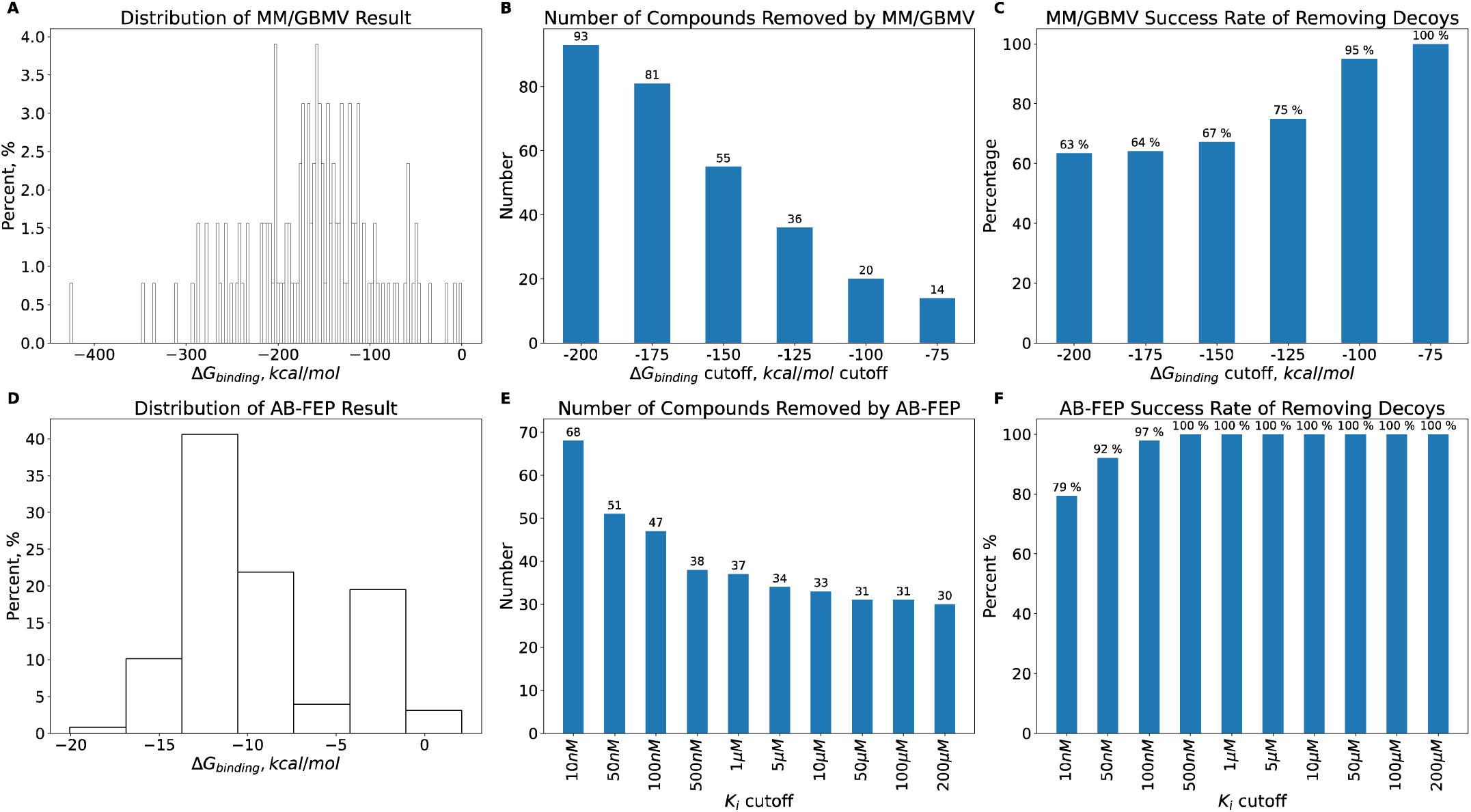
Re-scoring results for MM/GBMV: (A) Distribution of energy, (B) number of compounds removed and (C) success rate of removing decoys as a function of ΔG_binding_. The percentage is that of the compounds removed that are true “cheaters” at a given energy cut-off, with the remainder being false negatives; an ideal result is a maximum number of cheaters removed, and only cheaters removed (100%). Rescoring results for AB-FEP: (D) Distribution of energy, (E) number of compounds removed and (F) success rate of removing decoys as a function of K_d_. The percentage is as in panel C.

AB-FEP performed better. Here too, the range of energies was larger than the experimental values (Figure 8d), reaching affinities in the mid-femtomolar, and we considered two energy cutoffs to filter-out putative cheating molecules: molecules calculated by AB-FEP to have K_d_ values worse than 200 µM (essentially the experimental cutoff), and molecules with calculated K_d_ values worse than 1 µM . At the 200 µM cutoff, AB-FEP removed 30 of the 39 FACTS predicted cheating artifacts without flagging any false- negatives (true inhibitors) (Figure 8e). At the 1 µM cutoff it found 37 of the 39 cheating artifacts and classified another phenolate sulfonamide, a close analog of true inhibitors so arguably a plausible ligand, as a cheater, with no experimentally-confirmed false negatives found (Figure 8e). As AB-FEP cut-off K_d_ became more potent than 1 µM, the number of false negatives began to rise (Figure 8f). Detailed calculation results may be found in the Supplementary file (ampc-prospective- result.xlsx).

### Discussion and Conclusions

As our virtual libraries grow[2, 8, 19], both simulation [5] and experiment [Liu, 2024] suggest that docking results improve. Unfortunately, with this growth have emerged a small group of molecules that cheat our scoring functions and crowd the top scores of docking-ranked libraries [1, 5]. While rare, these molecules rise in sheer number as libraries grow; unchecked they may come to dominate the top-ranking molecules. A key observation from this study is that many of these scoring function “cheaters” may be recognized and deprioritized by an orthogonal scoring function. Perhaps the most compelling evidence for this comes from a prospective study against the model enzyme AmpC β-lactamase. Cross-filtering identified 39 “cheaters” from the very top-ranks of a docking campaign, and on experimental testing none bound substantially. Meanwhile, the method found another 89 compounds, often interspersed among the 39 “cheaters”, that were plausible ligands. On experimental testing, 57% of these 89 docking hits inhibited the enzyme. These results suggest that it is possible to identify molecules that cheat our scoring functions, filtering them out of top-ranking lists to reveal the more plausible and interesting high-ranking molecules.

Several caveats merit mentioning. Although we suspect that molecules that exploit scoring-function holes may be ubiquitous in large-library docking, we have only shown that they exist for DOCK3.7/3.8. Correspondingly, we have focused on physics-based scoring to cross-filter docking results, and while we might expect many methods to perform well in this role, we haven’t shown that.

These caveats should not obscure the key points of this study. We expect ever greater numbers of molecules to find holes in our scoring functions as docking libraries continue to grow and diversify[5]. Rescoring top-ranking molecules with a second scoring function that, whatever its own holes, is unlikely to share those of the primary scoring function, can help to eliminate these molecules, revealing the more interesting and plausible molecules that they rank among. This strategy thus may be generally useful in the field, and multiple scoring functions may be useful for rescoring. The FACTS method used here may be particularly well-suited to physics-based approaches, accordingly we provide easy-to-use rescoring and analysis scripts in the supplementary files.

## METHODS

Computational Details Unless otherwise specified, Open Babel [29] was used to generate random ligand conformations while ParamChem [30, 31] was used to prepare ligand topology and parameter files. The CHARMM C36 force fields [32] were used and molecular dynamics was performed in CHARMM [33]. The general AMBER force field (GAFF) [34] was used for comparison. Detailed molecular dynamic scripts are documented in the Supplementary Information.

Rescoring with implicit solvent model. We used the implicit solvent model FACTS (Fast Analytical Continuum Treatment of Solvation) model [20] and the GBMV (Generalized Born using Molecular Volume) [21] for rescoring. Unless otherwise noted, for each of the docked pose we performed a short minimization (1000 steps) using FACTS with CHARMM force field. The free energy at the unbound state is computed by first generating 20 random ligand conformations using the Open Babel functionality (obrotamer), followed by minimization.

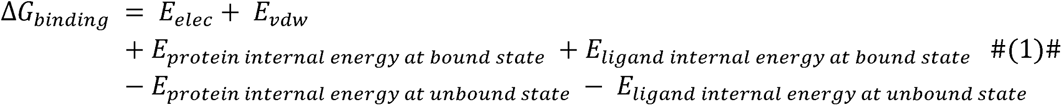

Cross-filtering with Bivariate Normal Distribution. For compounds with different topologies binding to the same receptor binding pocket, it is reasonable to assume these compounds have similar physio-chemical properties so that they could maintain similar key interactions with the binding pocket (i.e., normal distribution). However, the top ranking artifacts cheat one or multiple energy terms in the scoring function mainly because: (1) incorrect parameterization of the force field, and (2) the missing details in the scoring function. Therefore, using another docking method might help separating these artifacts.

As shown in Figure 3, one could draw an ellipse to determine the outliers. The width, height and the center of the ellipse are equivalent to the standard deviation and the mean of each of the normal distribution. While orientation of the ellipse can be determined by the covariance of the two variants. The size of the ellipse is defined by the user (i.e., σ cutoff). determined the ellipse based on the mean and standard deviation of each of the normal distribution and the covariance of the bivariate normal distribution The mathematical expression is eq. 1.

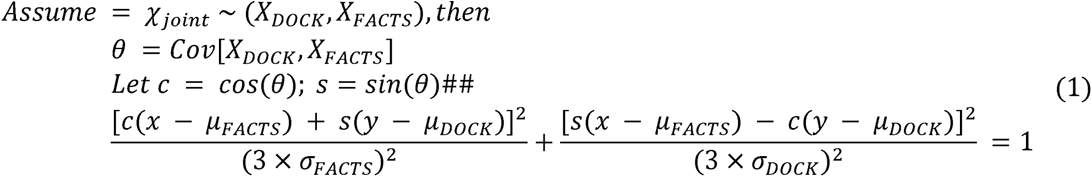

Calculating absolute binding free energy. Absolute Binding Free Energy (AB-FEP) calculations were performed with FEP+ on Schrodinger Web Services. The OPLS4 force field was used along with Force Field Builder, where missing parameters were calculated. The total MD simulation time for each ligand’s AB-FEP calculation was set to 1 ns. The docking poses used as the input for FEP were WScore docking results. All calculations are performed using Schrödinger Software Suite 2023-4 [35].

The MM/GBMV experiments were run up as follows [36]: for each compound, we collected top 5 docking poses. The bound state was minimized in vacuum with a maximum of 2000 steps before running molecular dynamics with the GBMV implicit solvent model. Hydrogen mass repartitioning was applied to reduce the computational cost. The total simulation length was 30 ns, and the last 1 ns of the simulation was used for the MM/GBMV calculation. Each simulation was repeated five times and the average energy computed. The lowest energy value among the five docking poses was considered the free energy of the bound state for that compound. The free energy for the ligand unbound state was computed with 25 random conformations generated by Open Babel, followed by the same MM/GBMV protocol.

AmpC β-lactamase enzymology. AmpC was purified as described.[37] All candidate inhibitors were dissolved in DMSO at 20 mM, and more dilute DMSO stocks were prepared as necessary so that the concentration of DMSO was held constant at 1% v/v in 50mM sodium cacodylate buffer, pH 6.5. AmpC activity and inhibition was monitored spectrophotometrically using either CENTA or nitrocefin as substrates. All assays included 0.01% Triton X-100 to reduce compound aggregation artifacts. Active compounds were further investigated for aggregation by dynamic light scattering (DLS) and by detergent-dependent inhibition of the counter-screening enzyme malate dehydrogenase.

For initial screening, the docking hits were diluted such that final concentrations in the reaction buffer was 200 µM, 100 µM and 40 µM. In these assays, the AmpC substrate nitrocefin^16^ was used, with an [S]/Km ratio of 0.56 (Km nitrocefin 180 µM; [S] = 100 µM) and 0.16 ([S] = 28 µM). The colorimetric assay was carried out using a BMG Labtech CLARIOstar for kinetic measurements of 50 seconds in 96-well format. IC50 values reflect the percentage inhibition fit to a dose-response equation in GraphPad Prism with a Hill coefficient set to one 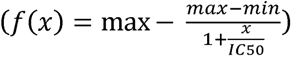. The K_i_ was calculated using the Cheng-Prusoff equation 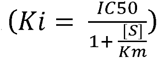 For eight of the most potent compounds, based on the initial three concentration-point results, full dose response curves were measured

### Malate Dehydrogenase Enzyme Inhibition Assay

Compounds were diluted to 100 µM in 50 mM KPi buffer, pH 7, at a final concentration of 1% DMSO (v/v). Samples were incubated with malate dehydrogenase (MDH) (Sigma, 442610) for 5 minutes. The reaction was initiated by adding 200 µM oxaloacetic acid (Sigma, 04126) and 200 µM nicotinamide adenine dinucleotide (NADH) (Sigma Aldrich,10128023001). Reaction was monitored for 80 seconds at an absorbance of 340 nm. Sample rates were divided by DMSO control rates. Compounds that displayed less than 35% inhibition were not considered inhibitors. Compounds that surpassed the enzyme inhibition threshold were screened as a concentration-response from 100 µM to 0.1 µM. Identified inhibitors were re-screened at 100µM in the presence of 0.01% (v/v) Triton-X 100 to determine detergent reversibility. All samples were screened in triplicate. Data were analyzed using GraphPad Prism version 10.2.3 (Boston, MA).

Dynamic Light Scattering. Compounds were diluted in filtered 50 mM KPi buffer, pH 7, at a final concentration of 1% DMSO (v/v). All compounds were initially screened at a top concentration of 100 µM using a DynaPro Plate Reader III. Samples that had a scattering intensity >1 X 10^7^ cnts/s scattering in this instrument were considered to form colloid-like particles and were re-screened as a concentration-response in eight-point half-log dilutions. Data were separated into two groups: aggregating concentrations (scattering > 1 X 10^7^cnts/s) and non-aggregating concentrations (scattering < 1 X 10^7^ cnts/s). A line was generated for each group and the point of intersection of the two lines serves as the critical aggregation concentration (CAC). All samples were screened in triplicate. Data were analyzed using GraphPad Prism.

## ASSOCIATED CONTENT

### Supporting information

Supporting Information is available free of charge on the ACS Publication website.

- Detailed description of bivariate distribution method development, filtering results with different implicit solvent models and force field parameters, DLS and Malate Dehydrogenase Enzyme Inhibition Assay results for top 10 compounds (PDF).
- Measured experimental K_i_ , AB-FEP, MM/GBMV and filtering results for all 128

prospectively tested compounds (xlsx).

- CHARMM and python scripts for cross-filtering, dataset for reproducing AmpC retrospective result (Figure 3) (tar.gz).

## Author information

Corresponding Author

Brian K. Shoichet - Department of Pharmaceutical Chemistry, University of California, San Francisco, 94158, United States.

Authors:

Yujin Wu - Department of Pharmaceutical Chemistry, University of California, San Francisco, 94158, United States.

Fangyu Liu - Department of Pharmaceutical Chemistry, University of California, San Francisco, 94158, United States.

Isabella Glenn - Department of Pharmaceutical Chemistry, University of California, San Francisco, 94158, United States.

Karla Fonseca-Valencia - Department of Pharmaceutical Chemistry, University of California, San Francisco, 94158, United States.

Lu Paris - Department of Pharmaceutical Chemistry, University of California, San Francisco, 94158, United States.

Yuyue Xiong – Schrödinger, Inc., 9868 Scranton Road, San Diego, California 92121, United States

Steven V. Jerome – Schrödinger, Inc., 1540 Broadway New York, New York 10036, United States

Charles L. Brooks III - Biophysics Program, University of Michigan, Ann Arbor, Michigan 48109, United States.

## Author Contributions

Y.W. developed cross-filtering methods. Y.W. and C.L.B. designed the rescoring setup with different implicit solvent models and MM/GBMV. Y.X. and S.J. performed AB-FEP calculations. Y.W. and F.L. identified the AmpC candidates for prospective study. F.L.,

I.S.G. and K.F.V. measured the experimental Ki against AmpC. L.P. performed colloidal aggregation screening using counter-screen enzymes MDH and DLS. B.K.S. supervised the project, designed experiments, and reviewed data. B.K.S. and Y.W. wrote the paper with input from the other authors; B.K.S. conceived the project.

## Notes

The authors declare the following competing financial interest(s): BKS is co-founder of BlueDolphin LLC, Epiodyne Inc, and Deep Apple Therapeutics, Inc., and serves on the SRB of Genentech, the SAB of Schrodinger LLC, and the SAB of Vilya Therapeutics. No other authors declare competing interests.

## Abbreviation Used

GAFF: general AMBER force field
FACTS: Fast Analytical Continuum Treatment of Solvation
FEP: free energy perturbation
AB-FEP: absolute binding-free energy perturbation
GBMV: Generalized Born using Molecular Volume
MM/GBMV: molecular mechanics/Generalized Born Molecular Volume
CRC: concentration-response curve

## Supporting information

Ampc prospective experimental and computational results.

Example cases for running cross-filtering

