## Supplementary figures and tables. for "Identifying Artifacts from Large Library Docking"

**Contents of SI:**

1. Table showing different cross-filtering setup.
2. Table showing experimental $K_{i}$ values calculated with 3 points and full CRC curve.
3. Figure showing joint distribution of solvation free energy using GBMV implicit solvent model.
4. Figure showing joint distribution of solvation free energy using FACTS implicit solvent model without minimization.
5. Figure showing joint distribution of solvation free energy using FACTS implicit solvent model with GAFF force field.
6. Figure showing percentage of compounds removed as a function of compound library screened.
7. Figure showing malate dehydrogenase enzyme (MDH) inhibition assay result.
8. DLS critical aggregation concentration assay for compound ZINCoG000003TUTt.

**Table S1** Cross-filtering Set up

| Implicit Solvent Model | Small Molecule Force Field Parameters | Minimization |
| --- | --- | --- |
| FACTSa | ParamChem | 1000 steps |
| FACTS | ParamChem | 0 steps |
| FACTS | GAFF | 1000 steps |
| GBMV | ParamChem | 1000 steps |

a Default setup in the proposed cross-filtering method.

Table S2: Experimental $K_{i}$ values calculated with 3 points and full CRC curve

| Receptor | $K_{i}$ (µM), 3 points | $K_{i}$ (µM), full CRC curve |
| --- | --- | --- |
| ZINCpH000009MUiv | 1.11 | 1.77 |
| ZINCpH000009MUiu | 6.24 | 2.44 |
| ZINCqo00000kNqT1 | 5.4 | 4.74 |
| ZINCpv0000008Z5r | 4.15 | 4.79 |
| ZINCpm00000drQg4 | 5.61 | 5.3 |
| ZINCoH000005YvzS | 6.02 | 5.41 |
| ZINCoL00000cODhJ | 14.6 | 13.16 |
| ZINCon00000986Sb | 12.57 | 15.19 |

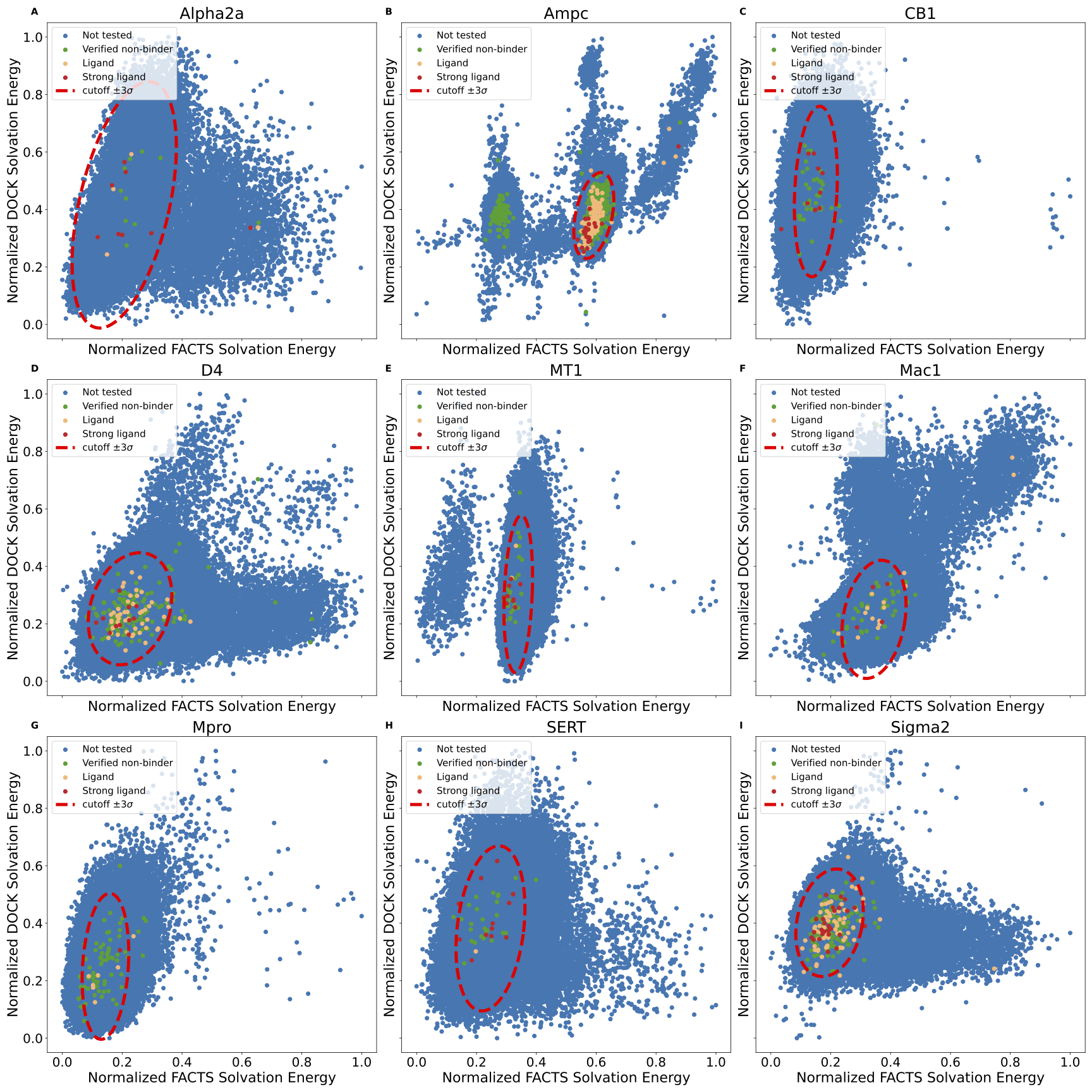

**Figure S1**: The joint distribution of the solvation free energy for receptor target **(A)** Alpha2a, **(B)** AmpC, **(C)** CB1, **(D)** D4, **(E)** MT1, **(F)** Mac1, **(G)** Mpro, **(H)** SERT and **(I)** Sigma2 using GBMV implicit solvent model. Potent ligands are colored red, and the rest of ligands are colored in orange. Experimentally validated non-binders and non-tested compounds are considered as decoys and colored blue. The true binders are color coded based on the binding affinity.

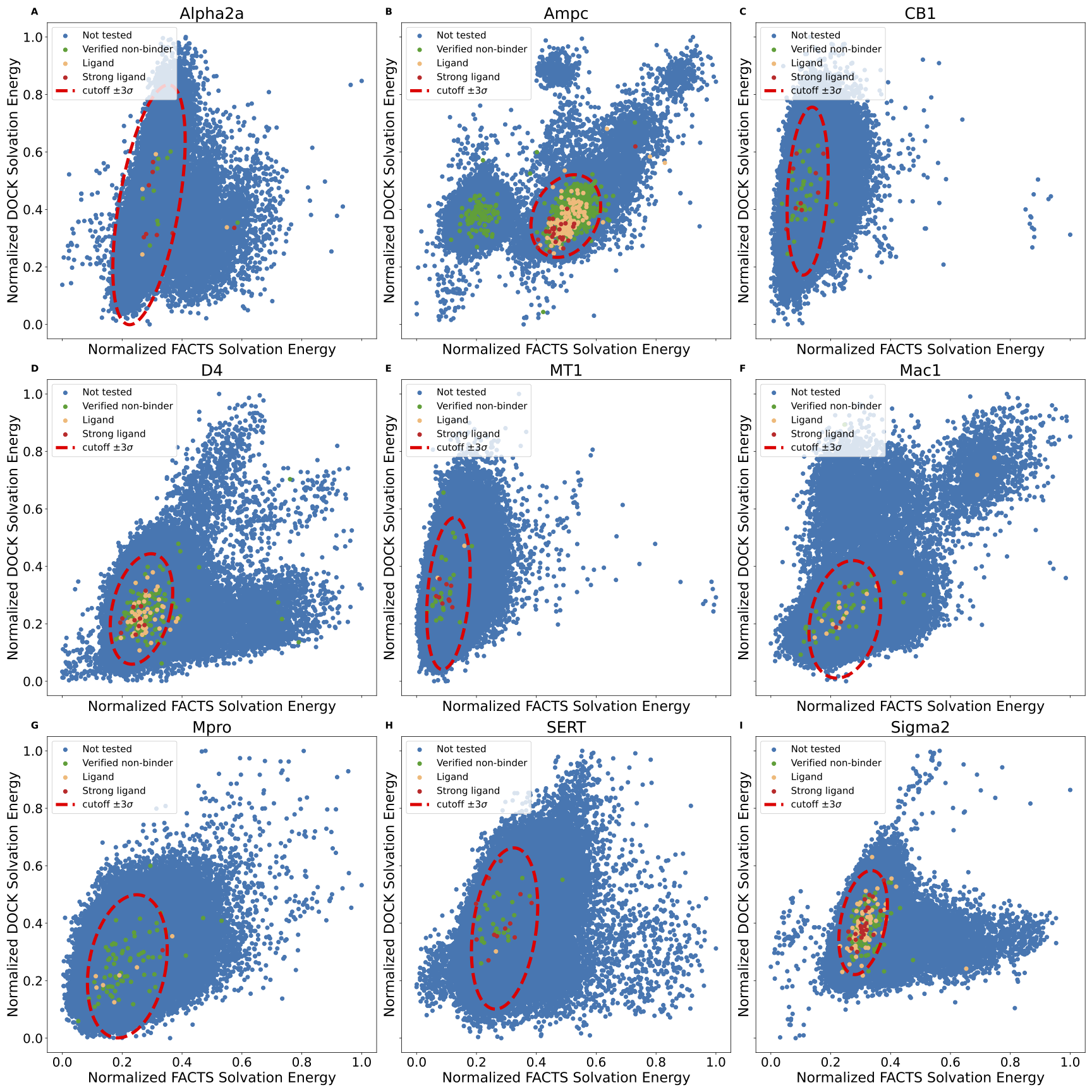

**Figure S2**: The joint distribution of the solvation free energy for receptor target **(A)** Alpha2a, **(B)** AmpC, **(C)** CB1, **(D)** D4, **(E)** MT1, **(F)** Mac1, **(G)** Mpro, **(H)** SERT and **(I)** Sigma2 using FACTS implicit solvent model without minimization. Potent ligands are colored red, and the rest of ligands are colored in orange. Experimentally validated non-binders and non-tested compounds are considered as decoys and colored blue. The true binders are color coded based on the binding affinity.

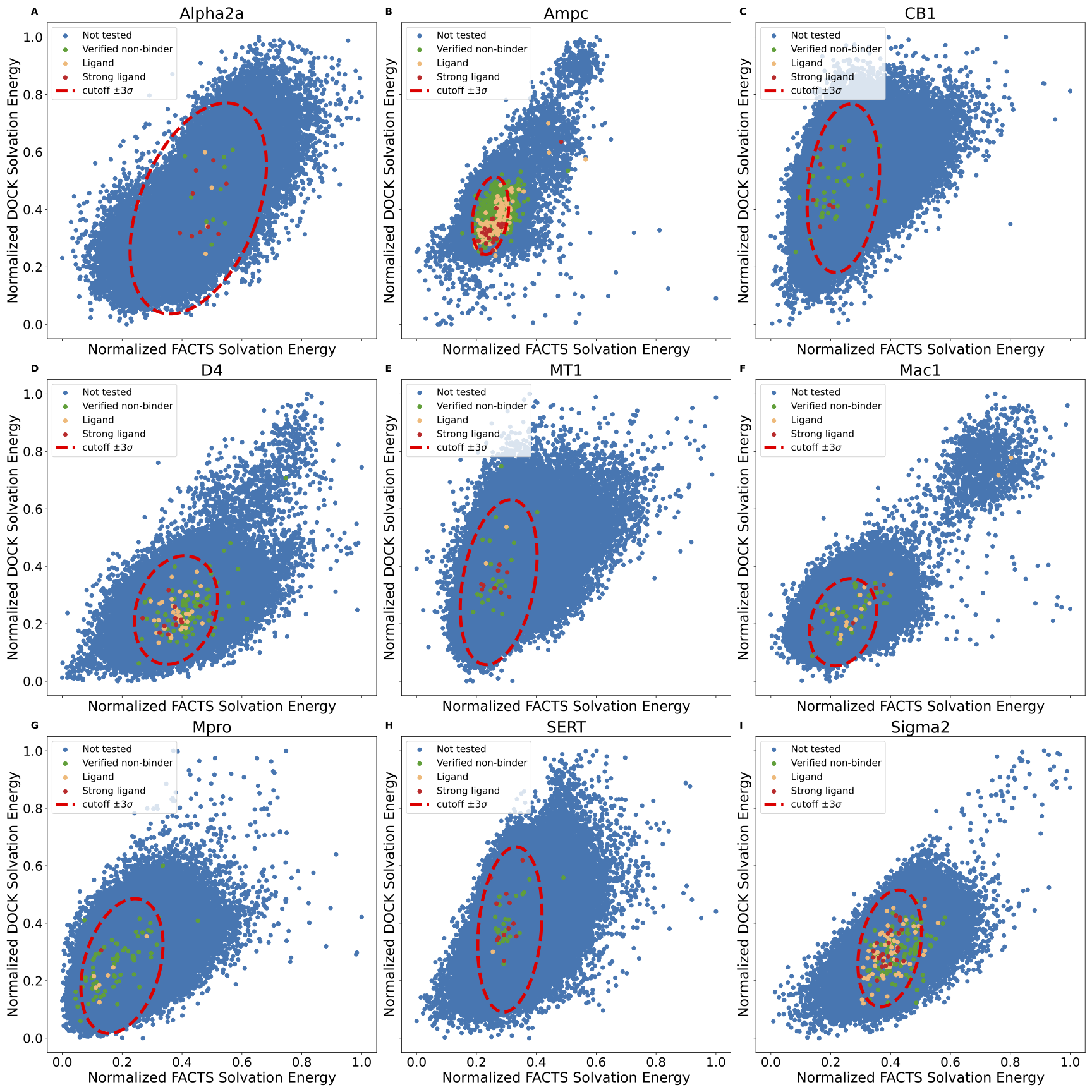

**Figure S3**: The joint distribution of the solvation free energy for receptor target **(A)** Alpha2a, **(B)** AmpC, **(C)** CB1, **(D)** D4, **(E)** MT1, **(F)** Mac1, **(G)** Mpro, **(H)** SERT and **(I)** Sigma2 using FACTS implicit solvent with GAFF force field. Potent ligands are colored red, and the rest of ligands are colored in orange. Experimentally validated non-binders and non-tested compounds are considered as decoys and colored blue. The true binders are color coded based on the binding affinity.

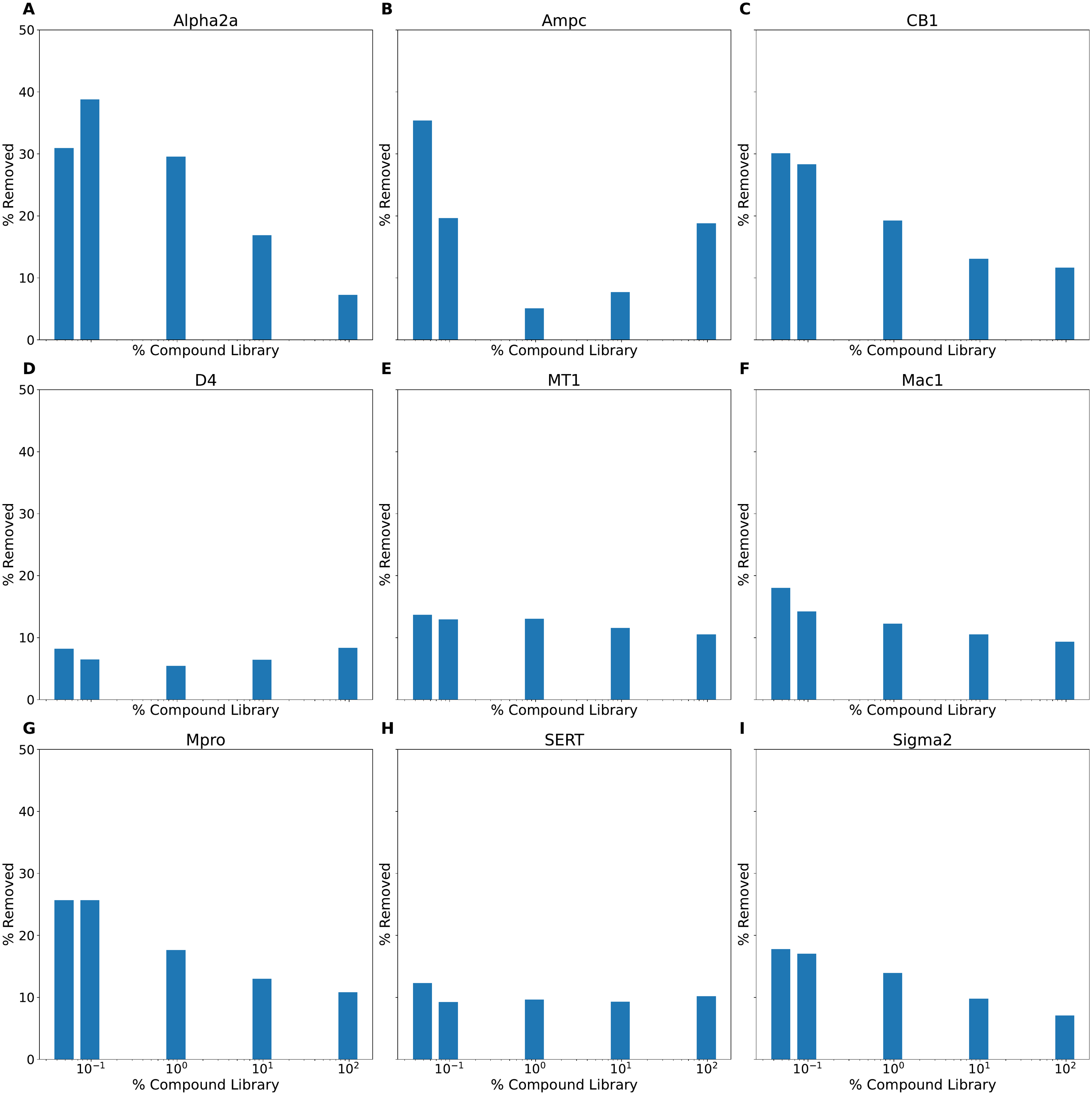

**Figure S4:** Percentage of compounds removed as a function of compound library screened (log scale) for receptor target **(A)** Alpha2a, **(B)** AmpC, **(C)** CB1, **(D)** D4, **(E)** MT1, **(F)** Mac1, **(G)** Mpro, **(H)** SERT and **(I)** Sigma2.

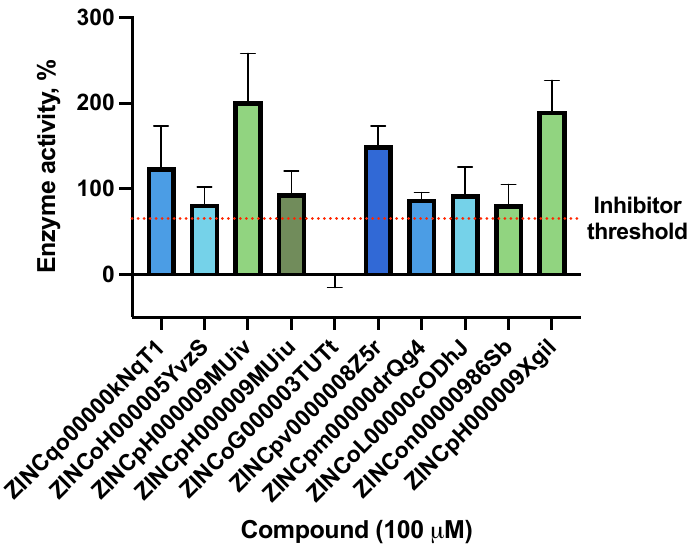

**Figure S5**: Malate dehydrogenase enzyme (MDH) inhibition assay. Compounds are tested at 100 µM and inhibitor cutoff is set at 65% MDH activity.

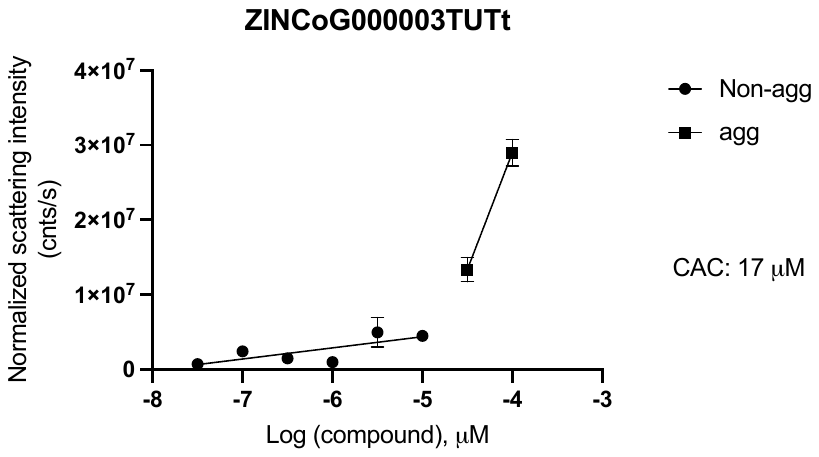

**Figure S6**: DLS critical aggregation concentration assay for compound ZINCoG000003TUTt.
