## Supplementary material for "Identifying Artifacts from Large Library Docking": Example cases for running cross-filtering: analysis_facts.py

```
## Import modules
import numpy as np
import pandas as pd
import seaborn as sns
import matplotlib as mpl
import matplotlib.pyplot as plt
from alive_progress import alive_bar
np.set_printoptions(suppress = True)


################################################################
## This is used to check the desolvation energy change 
## Compare with DOCK scoring function
## Plot bivariate distribution 
################################################################


################################################################
## Combine rescoring method and DOCK solvation free energy 
################################################################

## Loop through datasets and reorganize the docking results 
def calc_ener(hitList, dockResult) :
	'''
	------------------------------------------------
	facts_result/${compound}.dat -->
	each element in the result array is:
	index	meaning
	0	protein-ligand solvation contribution
	1 - 21	ligand solvation energy at unbound state
	------------------------------------------------
	Inputs :
	hitList		--> list of compounds that binds
		index	meaning
		0	ZINCID
		1	experimental Ki, uM
	dockResult	--> DOCK score
		index	meaning
		0	ZINCID	
		1	ligand polar
		2	ligand apolar
		3	DOCK score
	------------------------------------------------
	Returns :
	summary		--> summarized results 
	------------------------------------------------
	'''
	exp_ki = []
	ener_sol = []
	compound_Type = []
	with alive_bar(np.shape(dockResult)[0]) as bar :
		for idx in np.arange(np.shape(dockResult)[0]) :
			skip_this = False
			compound = dockResult[idx, 0]
			compound_Type.append("Ligand") if compound in hitList[:, 0] else compound_Type.append("Decoy")
			exp_ki.append(hitList[np.where(hitList[:, 0] == compound)[0], 1][0]) if compound in hitList[:, 0] else exp_ki.append(-1)
	
			try :
				facts_result = np.loadtxt("facts_result/{}.dat".format(compound), dtype = float)
				if np.sum(np.isnan(facts_result)) > 0 : skip_this = True
			except FileNotFoundError :
				skip_this = True
	
			if not skip_this : 
				l_sol = facts_result[1:]
				prob = l_sol - np.amin(l_sol)
				prob = - (1 / 0.593) * prob
				prob = np.exp(prob)
				prob = prob / np.sum(prob)
				l_sol = np.sum(prob * l_sol)
	
				pl_sol = facts_result[0]
				ener_sol.append(pl_sol - l_sol)
			else :
				ener_sol.append(np.inf)
			bar()

	ener_sol = np.asarray(ener_sol)
	ener_sol = ener_sol - np.amin(ener_sol)
	dock_sol = dockResult[:, 1].astype(float) + dockResult[:, 2].astype(float) 
	dock_sol = dock_sol - np.amin(dock_sol)

	summary = pd.DataFrame()
	summary["compound_Name"] = dockResult[:, 0]
	summary["compound_Type"] = compound_Type
	summary["DOCK_Score"] = dockResult[:, 3]
	summary["DOCK_Sol"] = dock_sol
	summary["FACTS_Sol"] = ener_sol
	summary["Ki, uM"] = np.array(exp_ki).astype(float)

	summary = summary.sort_values(by = ["DOCK_Score"], ignore_index = True, ascending = False)
	summary = summary.drop(summary[summary.FACTS_Sol == np.inf].index)
	summary = summary.drop(summary[summary.FACTS_Sol == np.nan].index)
	summary = summary.reset_index(drop = True)

	return summary.round(4)

################################################################
## Helper function for generate ellipse 
################################################################

## 2D Rotation matrix
def rot_2d(theta, data) :
	theta = np.radians(theta)
	c = np.cos(theta)
	s = np.sin(theta)
	R = np.asarray([[c, -s], [s, c]])
	data = np.dot(R, data.T)
	return data.T

## Generate random ellipse
def gen_ellipse(a, b, u, v, theta = 0) :
	'''		
	theta in degree unit
			|
			| b
		(u, v)	|
	----------------|----------------
	 		|        a
			|
			|
	'''
	point = np.linspace(0, 2 * np.pi, 1000)
	ellipse = np.zeros((len(point), 2))
	ellipse[:, 0] = a * np.cos(point) 
	ellipse[:, 1] = b * np.sin(point)
	rot_ellipse = rot_2d(theta, ellipse)
	
	ellipse[:, 0] = ellipse[:, 0] + u
	ellipse[:, 1] = ellipse[:, 1] + v

	rot_ellipse[:, 0] = rot_ellipse[:, 0] + u
	rot_ellipse[:, 1] = rot_ellipse[:, 1] + v

	return ellipse, rot_ellipse

## Check if point is in the ellipse
def check_within_ellipse(data, a, b, u, v, theta = 0) :
	theta = np.radians(theta)
	c = np.cos(theta)
	s = np.sin(theta)

	x = data[:, 0] - u
	y = data[:, 1] - v
	
	x_dist = (c * x + s * y) ** 2
	y_dist = (s * x - c * y) ** 2
	distance = x_dist / a ** 2 + y_dist / b ** 2

	return distance <= 1
	
################################################################
## Define 2D bivariate comparison plotting function 
################################################################

def facts_prediction(data, sigma = 3, filename = "facts_prediction", x = "FACTS_Sol", y = "DOCK_Sol") :
	'''
	Set up figure and subplots
	'''
	fig, axes = plt.subplots(3, 3, figsize = (18, 18))
	gs = axes[0, 0].get_gridspec()
	for idxI in [0, 1, 2] :
		for idxJ in [0, 1, 2] : axes[idxI, idxJ].remove()
	axX = fig.add_subplot(gs[2, 0:2])
	axY = fig.add_subplot(gs[0:2, 2])
	axSmall = fig.add_subplot(gs[2, 2])
	axBig = fig.add_subplot(gs[0:2, 0:2])

	dock = data[str(x)].to_numpy()
	ener = data[str(y)].to_numpy()
	data[str(x)] = dock / np.amax(dock)
	data[str(y)] = ener/ np.amax(ener)

	xrange = data[str(x)].to_numpy()
	xmin = np.floor(np.amin(xrange)) 
	xmax = np.ceil(np.amax(xrange))
	delta = 0.05 * (xmax - xmin)
	xmin = xmin - delta
	xmax = xmax + delta

	yrange = data[str(y)].to_numpy()
	ymin = np.floor(np.amin(yrange)) 
	ymax = np.ceil(np.amax(yrange))
	delta = 0.05 * (ymax - ymin)
	ymin = ymin - delta
	ymax = ymax + delta

	'''
	Plot axBig	--> normalized FACTS/DOCK solvation energy of each compound 
	'''
	label = np.unique(data["compound_Type"].to_numpy())
	cmap = mpl.colormaps["coolwarm"]
	tmpdf = data[data["compound_Type"] == label[0]]
	xPoint = tmpdf[str(x)].to_numpy()
	yPoint = tmpdf[str(y)].to_numpy()
	axBig.scatter(xPoint, yPoint, c = cmap(np.zeros(len(xPoint))), label = label[0])

	tmplig = data[data["compound_Type"] == label[1]]
	tmpdf = tmplig[tmplig["Ki, uM"] > 1000]
	xPoint = tmpdf[str(x)].to_numpy()
	yPoint = tmpdf[str(y)].to_numpy()
	if len(xPoint) > 0 : 
		axBig.scatter(xPoint, yPoint, c = cmap(np.ones(len(xPoint)) / 8 * 2), 
		label = label[1] + r": $ Ki > 1 mM$")

	tmplig = data[data["compound_Type"] == label[1]]
	tmpdf = tmplig[(tmplig["Ki, uM"] > 100) & (tmplig["Ki, uM"] <= 1000)]
	xPoint = tmpdf[str(x)].to_numpy()
	yPoint = tmpdf[str(y)].to_numpy()
	if len(xPoint) > 0 : 
		axBig.scatter(xPoint, yPoint, c = cmap(np.ones(len(xPoint)) / 8 * 3), 
		label = label[1] + r": $ 100 \mu M < Ki \leq 1 mM$")

	tmpdf = tmplig[(tmplig["Ki, uM"] <= 100) & (tmplig["Ki, uM"] > 10)]
	xPoint = tmpdf[str(x)].to_numpy()
	yPoint = tmpdf[str(y)].to_numpy()
	if len(xPoint) > 0 : 
		axBig.scatter(xPoint, yPoint, c = cmap(np.ones(len(xPoint)) / 8 * 4), 
		label = label[1] + r": $ 10 \mu M < Ki \leq 100 \mu M$")

	tmpdf = tmplig[(tmplig["Ki, uM"] <= 10) & (tmplig["Ki, uM"] > 1)]
	xPoint = tmpdf[str(x)].to_numpy()
	yPoint = tmpdf[str(y)].to_numpy()
	if len(xPoint) > 0 : 
		axBig.scatter(xPoint, yPoint, c = cmap(np.ones(len(xPoint)) / 8 * 5), 
		label = label[1] + r": $ 1 \mu M < Ki \leq 10 \mu M$")

	tmpdf = tmplig[(tmplig["Ki, uM"] <= 1) & (tmplig["Ki, uM"] > 0.1)]
	xPoint = tmpdf[str(x)].to_numpy()
	yPoint = tmpdf[str(y)].to_numpy()
	if len(xPoint) > 0 : 
		axBig.scatter(xPoint, yPoint, c = cmap(np.ones(len(xPoint)) / 8 * 6), 
		label = label[1] + r": $ 100 nM < Ki \leq 1 \mu M$")

	tmpdf = tmplig[(tmplig["Ki, uM"] <= 0.1) & (tmplig["Ki, uM"] > 0.01)]
	xPoint = tmpdf[str(x)].to_numpy()
	yPoint = tmpdf[str(y)].to_numpy()
	if len(xPoint) > 0 : 
		axBig.scatter(xPoint, yPoint, c = cmap(np.ones(len(xPoint)) / 8 * 7), 
		label = label[1] + r": $ 10 nM < Ki \leq 100 nM$")

	tmpdf = tmplig[tmplig["Ki, uM"] <= 0.01]
	xPoint = tmpdf[str(x)].to_numpy()
	yPoint = tmpdf[str(y)].to_numpy()
	if len(xPoint) > 0 : 
		axBig.scatter(xPoint, yPoint, c = cmap(np.ones(len(xPoint))), 
		label = label[1] + r": $Ki \leq 10 nM$")

	axBig.legend(fontsize = 15, loc = "upper left")
	
	'''
	Plot axBig	--> predicted ellipse 
	'''
	xy = data[[str(x), str(y)]].to_numpy()
	xPoint = xy[:, 0] 
	x_std = np.std(xPoint)
	x_mean = np.mean(xPoint)
	xy = xy[(xy[:, 0] < x_mean + 1 * x_std) * (xy[:, 0] > x_mean - 1 * x_std)]

	xPoint = xy[:, 0] 
	yPoint = xy[:, 1]
	x_std = np.std(xPoint)
	y_std = np.std(yPoint)
	x_mean = np.mean(xPoint)
	y_mean = np.mean(yPoint)

	cov = np.cov(xy.T)
	w, v = np.linalg.eig(cov)
	axis = v[0, :]
	theta = - np.degrees(np.arctan(axis[1] / axis[0]))
	ellipse, rot_ellipse = gen_ellipse(x_std * sigma, y_std * sigma, x_mean, y_mean ,theta)
	filter_result = check_within_ellipse(data[[str(x), str(y)]].to_numpy(), x_std * sigma, y_std * sigma, x_mean, y_mean, theta) 

	xy = data[[str(x), str(y)]].to_numpy()
	xy = xy[np.logical_not(filter_result)]
	axBig.plot(rot_ellipse[:, 0], rot_ellipse[:, 1], color = "red", ls = "dashed", lw = 2.5, label = r"cutoff $\pm$" + str(sigma) + r"$\sigma$")
	axBig.legend(fontsize = 15, loc = "upper left")

	axBig.set_xticklabels([])
	axBig.set_xlim([xmin, xmax])
	axBig.set_ylim([ymin, ymax])
	axBig.tick_params(axis = 'both', labelsize = 15)	
	axBig.set_ylabel('Normalized DOCK Solvation Energy', fontsize = 20)
	axBig.text(-0.075, 1.025, "A", size = 20, weight = "bold", transform = axBig.transAxes)
	axBig.set_title("Joint Distribution", fontsize = 25)

	'''
	Plot axX	--> Distribution of x-axis
	'''
	sns.kdeplot(data = data, x = x, ax = axX)
	axX.tick_params(axis = 'both', labelsize = 15)
	axX.legend([], [], frameon = False)
	axX.set_xlim([xmin, xmax])
	axX.set_ylabel('Percentage', fontsize = 20)
	axX.set_xlabel('Normalized FACTS Solvation Energy', fontsize = 20)
	axX.text(-0.075, 1.025, "C", size = 20, weight = "bold", transform = axX.transAxes)
	axX.set_title("Projected Distribution", fontsize = 25)

	'''
	Plot axY	--> Distribution of Y-axis
	'''
	sns.kdeplot(data = data, y = y, ax = axY)
	axY.tick_params(axis = 'both', labelsize = 15)
	axY.set_xlabel('Percentage', fontsize = 20)
	axY.legend([], [], frameon = False)
	axY.set_ylim([ymin, ymax])
	axY.set_yticklabels([])
	axY.set_ylabel('')
	axY.text(-0.15, 1.025, "B", size = 20, weight = "bold", transform = axY.transAxes)
	axY.set_title("Projected Distribution", fontsize = 25, rotation = "vertical", 
			va = "center", ha = "left", x = 1.025, y = 0.5) 

	'''
	Plot axSmall	--> Bar plot of removed compound percentage 
		D 	--> Data
		F	--> Pass filtering 
		R	--> Removed
	'''
	filtered = data.loc[filter_result]
	filtered = filtered.sort_values(by = ["DOCK_Score"], ignore_index = True)
	filtered.reset_index(drop = True, inplace = True)

	num_D = []
	num_F = []
	cutoff = np.array([0.1, 1, 10, 100])

	for idx in np.arange(len(cutoff) -1 ): 
		tmpdf = data[(data["Ki, uM"] <= cutoff[idx + 1]) & (data["Ki, uM"] > cutoff[idx])]
		num_D.append(tmpdf.shape[0])

		tmpdf = filtered[(filtered["Ki, uM"] <= cutoff[idx + 1]) & (filtered["Ki, uM"] > cutoff[idx])]
		num_F.append(tmpdf.shape[0])

	num_D.append(data[data["compound_Type"] == "Decoy"].shape[0])
	num_F.append(filtered[filtered["compound_Type"] == "Decoy"].shape[0])
	
	num_D = np.asarray(num_D)
	num_F = np.asarray(num_F)
	num_R = num_D - num_F

	bars = axSmall.bar(np.arange(len(num_R)), num_R, width = 0.3, label = "Number")
	for bar in bars :
		height = bar.get_height()
		axSmall.annotate("# {}".format(int(height)), xy = (bar.get_x() + bar.get_width() / 2, height), 
				xytext = (0, 3), textcoords = "offset points", ha = "center", va = "bottom")

	axSmall.tick_params(axis = 'both', labelsize = 15)	
	axSmall.set_ylabel("Percentage/Number", fontsize = 20)
	axSmall.set_xlabel("Compound", fontsize = 20)
	axSmall.set_xticks(np.arange(len(num_R)))
	axSmall.set_xticklabels([r"$100 nM$" "\n" r"$\sim$" "\n" r"$1 \mu M$", 
				 r"$1 \mu M$" "\n" r"$\sim$" "\n" r"$10 \mu M$",
				 r"$10 \mu M$" "\n" r"$\sim$" "\n" r"$100 \mu M$", "Decoy"])
	axSmall.text(-0.15, 1.025, "D", size = 20, weight = "bold", transform = axSmall.transAxes)
	axSmall.set_title("Compounds Filtered", fontsize = 25)
	axSmall.legend(fontsize = 15, loc = "upper left")

	plt.tight_layout(pad = 1.15)
	plt.savefig("{}.pdf".format(filename), dpi = 800)
	plt.close()

	'''
	Save filtering result
	'''
	predict = np.chararray(len(filter_result), itemsize = 20)
	predict[filter_result] = "Plausible Actives"
	predict[np.logical_not(filter_result)] = "Likely Cheater"
	data["Prediction"] = predict
	data.to_csv("{}.tsv".format(filename), sep = "\t")

	return data 

################################################################
## Compare bivariate distribution with different force field 
################################################################

dockResult = np.loadtxt("./dockResult", dtype = str)
hitList = np.loadtxt("./hitList", dtype = str)
summary = calc_ener(hitList, dockResult) 
data = facts_prediction(summary)
print(data)
```
